# Platelet endocytosis and α-granule cargo packaging are essential for normal skin wound healing

**DOI:** 10.1101/2025.02.01.636051

**Authors:** Daniëlle M. Coenen, Hammodah R. Alfar, Sidney W. Whiteheart

**Author notes:** Corresponding Author: Sidney W. Whiteheart, Ph.D. FAHA, Department of Molecular and Cellular Biochemistry University of Kentucky, College of Medicine, B361 BBSRB, 741 S Limestone, Lexington, KY 40508, USA.

## Abstract

The high prevalence of chronic wounds, *i.e.,* 2.5-3% of the US population, causes a large social and financial burden. Physiological wound healing is a multi-step process that involves different cell types and growth factors. Platelet-rich plasma or platelet-derived factors have been used to accelerate wound repair, but their use has been controversial with mixed results. Thus, a detailed functional understanding of platelet functions in wound healing beyond hemostasis is needed. This study investigated the importance of platelet α-granule cargo packaging and endocytosis in a dorsal full-thickness excisional skin wound model using mice with defects in α-granule cargo packaging (Nbeal2^-/-^ mice) and endocytosis (platelet-specific Arf6^-/-^ and VAMP2/3^Δ^ mice). We found that proper kinetic and morphological healing of dorsal skin wounds in mice requires both *de novo* as well as endocytosed platelet α-granule cargo. Histological and morphometric analyses of cross-sectional wound sections illustrated that mice with defects in α-granule cargo packaging or platelet endocytosis had delayed (epi)dermal regeneration in both earlier and advanced healing. This was reflected by reductions in wound collagen and muscle/keratin content, delayed scab formation and/or resolution, re-epithelialization, and cell migration and proliferation. Molecular profiling analysis of wound extracts showed that the impact of platelet function extends beyond hemostasis to the inflammation, proliferation, and tissue remodeling phases via altered expression of several bioactive molecules, including IL-1β, VEGF, MMP-9, and TIMP-1. These findings provide a basis for advances in clinical wound care through a better understanding of key mechanistic processes and cellular interactions in (patho)physiological wound healing.

**Key points:**

- *De novo* and endocytosed platelet α-granule cargo support physiological skin wound healing
- Platelet function in wound healing extends to the inflammation, proliferation, and tissue remodeling phases

## Introduction

Currently, around 2.5-3% of the US population (± 10.5 million people) suffer from chronic wounds, which is higher than the prevalence of heart failure.^1–4^ With a globally increasing obese and (pre-)diabetic populations, these numbers are expected to increase, causing a large social and financial burden; reducing quality of life, and placing pressure on healthcare and the economy.^3,5–7^ Even though total expenditures declined over the past years due to technological and medical improvements, (chronic) wound care still costs more than $22.5 billion per year. Wound dressings and gels are clinically used to treat acute and chronic wounds, frequently supplemented with plasma and/or cellular or molecular constituents, such as growth factors or cytokines, and often derived from platelets.^8–11^ However, their use has been controversial with mixed results, in part due to a lack of (experimental) standardization and underexplored mechanisms.^12–19^

The physiological process of (skin) wound healing can be divided into four phases, *i.e.,* hemostasis, inflammation, proliferation, and maturation or tissue remodeling.^20–24^ Directly after wounding, platelets accumulate at the site of injury to form a hemostatic plug to stop bleeding. This is quickly followed by the recruitment of neutrophils to elicit an inflammatory response, after which monocytes and macrophages break down excessive pathogenic components and clear the environment of residual debris. New tissue is formed during the proliferation phase, requiring multiple cell types including keratinocytes, macrophages, (myo)fibroblasts, and endothelial cells. The formation of these connective, or granulation, tissues is accompanied by revascularization/angiogenesis, synthesis of extracellular matrix proteins, wound contraction, and re-epithelialization. Final wound closure and resolution is realized by removal of cellular residue and matrix remodeling, through turnover of type III collagen into collagen type I to replace the granulation tissue with normal connective, or scar, tissue.^21,25–27^ Despite being separate processes and the predominance of specific cell types in each phase, there are overlap between the phases.^22,25^ Interactions between the various cell types and initiation of the different processes in wound healing are coordinated via the release and action of growth factors, chemokines, and other soluble molecules.^28^ The most common and important mediators in wound healing are EGF, FGF2, IGF, MMPs, PDGF, TGF-β, TIMPs, TNF-α, VEGF, and numerous interleukins. Throughout these different phases, such factors are released in a coordinated manner by several different cell types, including platelets.^25,28,29^ The cellular and molecular mechanisms of wound repair are described in detail in recent reviews from Gurtner *et al.*,^21^ Rodrigues *et al.*,^26^ Barrientos *et al.*,^29^ Qing,^30^ and Oscar Peña and Paul Martin.^22^ However, the specific roles of platelets in wound healing, beyond hemostasis, are less clear.

Platelets have a dominant role in maintaining vascular integrity and communicate (in)directly with their environment via SNARE-mediated, *i.e.,* granular, and unconventional, *i.e.,* non-granular, protein secretion.^31–34^ In platelets, granular cargo proteins are mainly secreted via α-granules, dense granules, and lysosomes.^32,35^ Dense granules contain a host of small molecules that are rapidly released and critical to primary hemostasis. Lysosomes have the slowest release response and contain acid hydrolases and proteases. Platelet α-granules are the most abundant and diverse; their cargo is not only synthesized and packaged in progenitor megakaryocytes (*de novo* cargo), but also derived from several endocytic pathways. Platelet α-granule cargo includes angiogenic factors, chemokines, coagulation factors, growth factors, immune mediators, and (inhibitors of) metalloproteinases (Supplemental Table 1 and Table 1).^36^ Given this wealth of potential effectors, it is remarkable that platelet function in wound healing, other than in the hemostasis phase, has been understudied. The wide range of chemokines, cytokines, and growth factors present in platelet α-granules and the variety of (patho)physiological processes regulated by these, including inflammation, angiogenesis, and tissue remodeling point to a pivotal role for platelets beyond initial bleeding cessation.^37–43^ Over the last decade, only a few studies have suggested that platelets and their granule cargo orchestrate physiological skin wound healing.^44–47^

**Table 1.** Cargo known to be endocytosed by megakaryocytes (MKs) and/or platelets.

| **Full name** | **Endocytosed**<br>**by platelets** | **Endocytosed**<br>**by MKs** | **References** |
| --- | --- | --- | --- |
| Endostatin | Yes | Yes | 98-100 |
| Factor V | No | Yes | 36,101 |
| Factor XIII | Yes | N.I. | 102,103 |
| Fibroblast growth factor 2 | Yes | N.I. | 99 |
| Fibrinogen | Yes | Yes | 36,98,104-109 |
| Insulin-like growth factor-1 | No | Yes | 36,110 |
| Platelet-derived growth factor | Yes | N.I. | 99 |
| Serum albumin | Yes | Yes | 36,107,108,111 |
| Vascular endothelial growth factor | Yes | N.I. | 36,67,99 |

MK, megakaryocyte, N.I, not identified.

In this study, we sought to better understand platelet’s role in wound healing using transgenic mouse models where specific aspects of platelet secretion have been manipulated, in particular α-granule cargo packaging and endocytosis. The Nbeal2^-/-^ mice have a defect in α-granule biogenesis and cargo packaging, such that their α-granules almost completely devoid of soluble cargo.^48–50^ Because of the multiple sources of the α-granule content, we also used mice with platelet-specific defects in endocytosis, *i.e.,* Arf6^-/-^ and VAMP2/3^Δ^ mice. The platelet-specific Arf6^-/-^ and VAMP2/3^Δ^ mice have decreased concentrations of endocytosed proteins, *e.g.,* fibrinogen, factor XIII, and vascular endothelial growth factor (VEGF).^36,51,52^ Utilizing these different mouse strains allowed us not only to conduct a more in-depth comparison of *de novo* vs. endocytosed α-granule cargo proteins, but also to explore different phases of endocytic trafficking as Arf6 is involved in initial cargo uptake,^36,51^ while deletion of VAMP2 and 3 affect subsequent trafficking of the endocytosed proteins.^32,53^ We found that alterations of the cargo packaging pathways greatly affected skin wound healing. Wounds from Nbeal2^-/-^, Arf6^-/-^, and VAMP2/3^Δ^ mice healed slower and morphologically and histologically differed from wildtype. (Epi)dermal regeneration was delayed with complications in scab formation and/or resolution, re-epithelialization, and cell migration and proliferation. We observed specific relevance of *de novo* vs. endocytosed α-granule cargo depending on the healing stage. The alterations in platelet α-granule cargo packaging resulted in changes in the expression of bioactive molecules in skin wound extracts, disturbing inflammation, proliferation, and tissue remodeling. Taken together, our data demonstrates an important role for platelet α-granule cargo packaging and platelet endocytosis in healing skin wounds that extend beyond initial hemostasis. Such insights contribute to potential improved treatments of both acute and chronic wounds.

## Methods

### Animals

Nbeal2 knockout (KO; Nbeal2^-/-^) mice were obtained from the Mutant Mouse Resource and Research Center (MMRRC) at the University of California at Davis.^54^ Coding exons 4 through 11 were targeted by homologous recombination.

Platelet-specific Arf6-deficient (Arf6^flox/flox^::PF4Cre^+^; Arf6^-/-^) mice were generated by crossing Arf6^flox/flox^ mice (generous gift of Dr. Y. Kanaho, University of Tsukuba, Tsukuba, Japan) with PF4-promoter-driven Cre-recombinase transgenic mice (kindly provided by Dr. R. Skoda, University Hospital, Basel, Switzerland).^55,56^

Platelet-specific VAMP2/3-deficient (VAMP2/3{ROSA::STOP^fl/fl^::TetTox::PF4Cre^+^}; VAMP2/3^Δ^) mice were generated by crossing RC::PFtox mice (generously provided by Dr. S. Dymecki, Harvard Medical School, Boston, MA) with the previously mentioned PF4-Cre mice.^52^

Age- and sex-matched C57BL/6J wildtype (WT) mice, purchased from Jackson Laboratory and bred in-house, were used as controls. All mice used were between 8 and 10 weeks old, with an even distribution of male and female mice. Mice were bred and maintained on pelleted paper bedding (Teklad 7084) under standard husbandry conditions in the animal facility of the University of Kentucky and fed regular chow diet *ad libitum*. All experiments performed were approved by the Institutional Animal Care and Use Committee (IACUC) of the University of Kentucky under protocol number 2021-3853. Genotyping strategies are described in Supplemental Methods.

### Full-thickness excisional skin wound model

Two dorsal full-thickness circular excisions were made on shaved mice with a 4-mm biopsy punch (Sklar, Merit™ Keyes, supplier no. 98-404) as described.^57^ Briefly, the dorsal skin was raised and folded at midline, after which the animal was placed in lateral position and the biopsy punch was pressed through the two skin layers to create two symmetrical full-thickness wounds. The wound area was measured using calipers, immediately after wounding and daily, for a maximum of seven days after wounding. The healing kinetics were determined by comparing the area of the healing wounds with the initial wound size and presented as percentage of the initial wound size.

### Skin histology and morphometric wound analysis

At the time of wounding, and three and seven days afterwards, skin tissue was collected and fixed overnight in 10% formalin. The tissues were processed and paraffin-embedded at the Pathology Research Core at the University of Kentucky, using standard protocols. Cross-sectional skin sections (5 μm) were used for hematoxylin and eosin (H&E) and Masson’s Trichrome staining and imaged at 20x using a Zeiss Axioscan Z7 automated slide scanner, available at the Light Microscopy Core at the University of Kentucky.

H&E-stained sections were manually analyzed for several wound morphometrics using visual scoring of the images and freehand line and spline contour tools in Zeiss Zen lite version 3.10, as shown in Supplemental Figure 1 and based on Wichaiyo *et al.*.^45^ Sections were scored based on the presence or absence of a scab. Providing its presence, scab area was measured (Supplemental Figure 1A). Furthermore, formation of a full or partial epidermal layer was assessed. Sections with a partially formed epidermal layer were further analyzed to calculate percentage re-epithelialization, or epidermal regeneration (Supplemental Figure 1B, C).^45,58,59^ Percentage re-epithelialization was calculated using the following formula: % re-epithelialization = [(E_1_+E_2_)/W_0_] × 100. E_1_ and E_2_ is the distance of epidermal regeneration on either side of the scab, whereas W_0_ is the total distance between the two wound edges. The wound edge is the area where the normal continuous skin epidermis transitions into a thickening keratinocyte layer. Last, the area of granulation tissue was measured (Supplemental Figure 1D). Granulation tissue consists primarily of fibroblasts, collagen, macrophages, and new blood vessels, and its presence is characteristic for the proliferative phase of wound healing.^45,60–63^

Muscle/keratin and collagen contents were determined in the Masson’s Trichrome-stained sections using Fiji ImageJ.^45,47^ Briefly, color deconvolution (Masson’s Trichrome) was used to separate muscle/keratin (red) and collagen (blue) from green and thresholding was applied using the Otsu method. The intensity of the blue and red color was normalized by the measured area and expressed as fold change relative to healthy skin tissue of wildtype mice.

### Molecular profiling analysis

Extensive molecular profiling analysis of skin wound extracts was performed using a Luminex Discovery Assay from R&D Systems. Healthy skin and three- and seven-day wound specimens were homogenized with an electric homogenizer in 300 μl ice-cold PBS containing 600 mM NaCl, 1x Halt^TM^ Protease Inhibitor Cocktail (Thermo Fisher Scientific), and 1x, or 5mM, ethylenediaminetetraacetic acid (EDTA; Thermo Fisher Scientific). The homogenizer was rinsed with 200 μl PBS, and the samples, in a total volume of 500 μl, were placed on ice and lysed with 1% Triton X-100. Every 15 min., the samples were briefly vortexed and after 30 min., the tissue was crushed with a disposable pestle and centrifuged at 17,000 g for 10 min. Supernatant was transferred to a new tube and both supernatant and homogenized tissue were stored at -80 °C for further analysis.

Mouse premixed multi-analyte Luminex Discovery Assays (R&D Systems) were used to measure the levels of several bioactive molecules in skin wound extracts (Supplemental Table 1) according to the manufacturer’s instructions, except for the sample dilutions. Healthy skin samples were diluted 1:2, while extracts from tissue collected three and seven days after wounding were measured after being diluted twice and after a 50-time dilution. Plates were read with a Luminex MAGPIX analyzer, collecting Median Fluorescence Intensity (MFI) of 50 counts/region in 50 μl sample volume.

Final concentrations of the Luminex-measured analytes were normalized against total protein levels in the tissue, determined with the Bicinchoninic acid (BCA) assay. Briefly, a protein standard was made with Bovine Serum Albumin Fraction V (Millipore Sigma). Samples were used undiluted and 10x or 20x diluted in sterile water. Reagent A (sodium carbonate, sodium bicarbonate, BCA^TM^ detection reagent, and sodium tartrate in 0.1N sodium hydroxide; Thermo Fischer Scientific) was mixed with reagent B (4% cupric sulfate) in a 50:1 ratio. 200 μl of the reagent A/B solution was added to 25 μl of standards and samples in a 96-well plate. After 20-30 min. of incubation at 37°C, absorbance was measured at 570 nm.

### Data analysis and statistics

Wound healing kinetics, wound histology, and wound morphometrics were analyzed with a two-way ANOVA with Šidák’s (WT vs. Nbeal2^-/-^) or Dunnett’s (WT vs. Arf6^-/-^ and VAMP2/3^Δ^) post-hoc testing to correct for multiple comparisons. Spearman correlation analyses were conducted on analyte levels in the wound extracts and wound resolution. All data was statistically analyzed with Graphpad Prism version 10.4.0 and shown as mean ± standard deviation (SD) or median ± interquartile range (IQR). Differences were considered significant with a p-value of less than 0.05.

### Data sharing statement

For original data, please contact.

## Results

### *De novo* and endocytosed platelet α-granule cargo support physiological skin wound healing

To better understand the role of platelet α-granule cargo, both *de novo* synthesized and endocytosed, in physiological skin wound healing, we applied a full-thickness excisional skin wounding model to wildtype mice (C57BL/6J) and mice with defects in α-granule cargo packaging (Nbeal2^-/-^) or endocytosis (Arf6^-/-^ and VAMP2/3^Δ^). In this model, dorsal skin wounds made on wildtype mice healed almost completely (mean = 85.6% of initial wound size, *n* = 15) in seven days (Figure 1). Scab formation occurred around day three post-wounding and its resolution took place after five to seven days (Figure 1A). Skin wound healing in Nbeal2^-/-^ mice was severely impaired as only 72.4% of the initial wound size was healed seven days post-wounding (p = 0.0127, *n* = 10; Figure 1A, B). Moreover, Nbeal2^-/-^ mice had a distinctive wound morphology as scab formation and scab resolution were considerably hampered (Figure 1A). Like Nbeal2^-/-^ mice, Arf6^-/-^ and VAMP2/3^Δ^ mice also presented with remarkably slower wound healing (a seven-day decrease of 79.2% (p = 0.2584, *n* = 16) and 75.3% (p = 0.0646, *n* = 11) of initial wound size, respectively; Figure 1A, C). Notably, healing kinetics of Arf6^-/-^ mice did not deviate from that of wildtype mice until the second day post-wounding. These data suggest that alterations in both *de novo* and endocytosed platelet α-granule cargo affect skin wound healing with particularly *de novo* proteins being pivotal in the initial healing phase.

**Figure 1.**
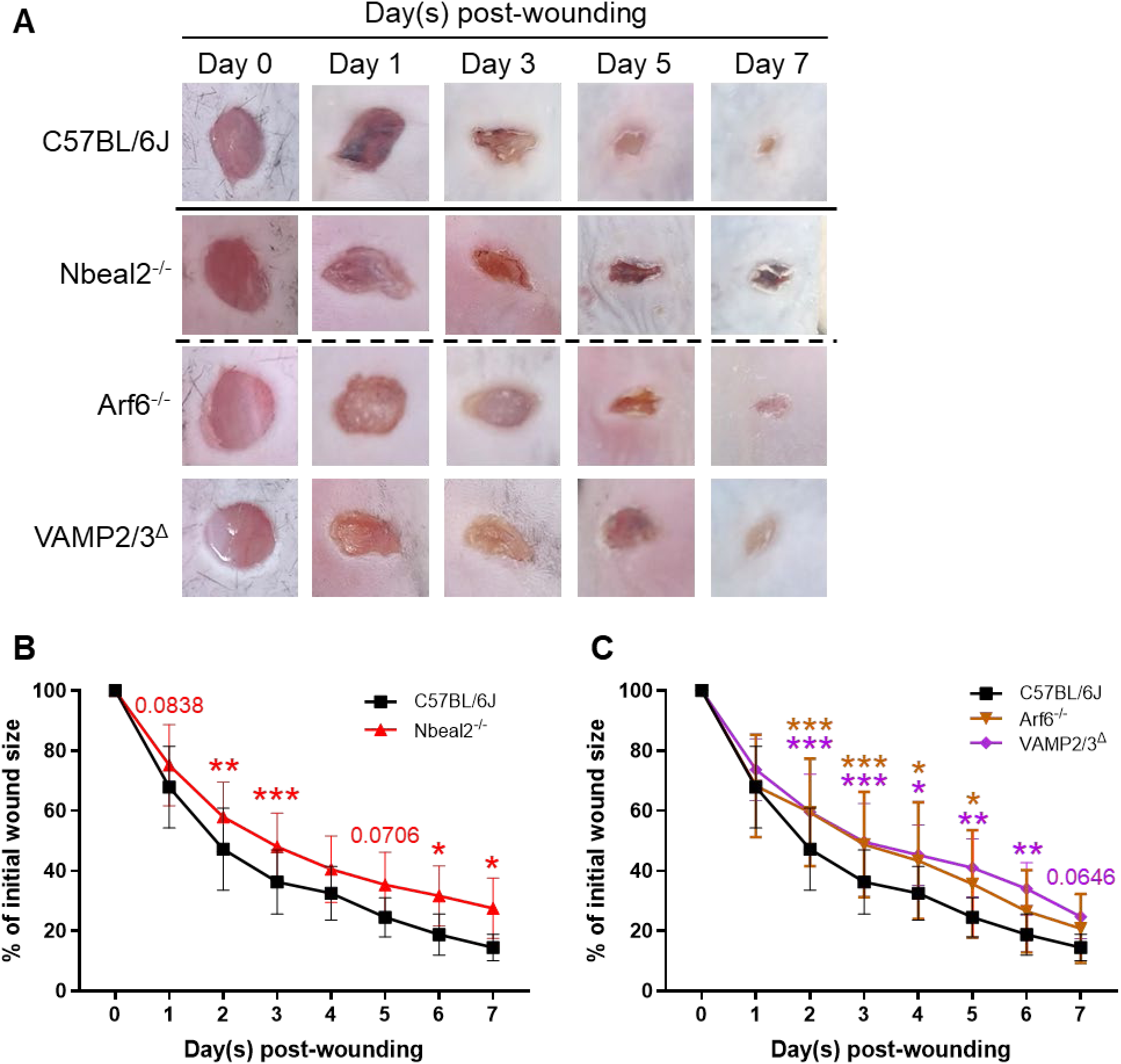
Skin wound healing is impaired in mice deficient in α-granule cargo packaging or endocytosis. **(A)** Representative macroscopic images of (healing) dorsal full-thickness excisional wounds on indicated days. **(B, C)** Percentage healing over time compared to baseline (day 0) in wildtype mice and mice with defects in α-granule cargo packaging **(B)** or endocytosis **(C)**. Mean ± S.D., *n* = 19-33 (day 0 – day 3), 10-16 (day 4 – day 7), * p < 0.05, ** p < 0.01, *** p < 0.001. Statistics: two-way ANOVA followed by Šidák’s **(B)** and Dunnett’s **(C)** multiple comparisons tests

### Disrupting α-granule cargo release and platelet endocytosis delays (epi)dermal regeneration of skin wounds

The full-thickness skin wound model affects all skin layers, *i.e.,* epidermis, dermis, and hypodermis, thus we first analyzed gross wound morphometrics of H&E-stained healthy skin and (healing) wounds (see Supplemental Figure 1 for definitions). In cross-sections of healthy biopsies, the complete skin structure could be identified, *e.g.,* hair follicles, glands, blood vessels, subcutaneous tissue, and the *panniculus carnosus* muscle (Figures 2A). Healthy skin from Nbeal2^-/-^, Arf6^-/-^, and VAMP2/3^Δ^ mice were histologically similar to wildtype (Figure 2A). Except for one wildtype animal, all mice developed a scab within three days (Figure 2A, B, D). In most of the wildtype and Arf6^-/-^ mice, the scab was resolved by seven days post-wounding (Figure 2A, B, D). At the same timepoint, wounds of all Nbeal2^-/-^ and the majority of the VAMP2/3^Δ^ mice still presented with a scab (Figure 2A, B, D). VAMP2/3^Δ^ and Nbeal2^-/-^ mice showed a trend in increased scab area three and seven days after wounding (p = 0.16 and p = 0.14, respectively; Figure 2B, D). Complications in scab formation and resolution were reflected by the lack of epidermal regeneration, as defined by the formation of a full or partial epidermal layer and, in case of the latter, percentage of re-epithelialization (Figure 2A, F-I). Epidermal regeneration started shortly after wounding, but after three days, none of the animals had a full epidermal layer as the epidermis was still obstructed by the presence of a large scab (Figure 2A, B, D, F, H). In wildtype mice, almost 60% of the initial wound was closed by a new epidermal layer (Figure 2G, I). Re-epithelialization was slightly reduced in Nbeal2^-/-^ (41%, p = 0.2442), Arf6^-/-^ (36%, p = 0.1970), and VAMP2/3^Δ^ (37%, p = 0.1893) mice. After seven days, most wildtype and Arf6^-/-^ mice had a complete epidermal layer, while the amount of epidermal re-epithelialization was still decreased in Nbeal2^-/-^ and VAMP2/3^Δ^ mice (p = 0.0074 and p = 0.1360; Figure 2F-I). A hallmark of skin wound healing is the formation of granulation tissue, which builds up the injured area with a foundation composed of newly recruited cells and developing blood vessels and is indicative of the proliferative phase of wound healing.^45,60–63^ We found a smaller area of granulation tissue in Nbeal2^-/-^ and Arf6^-/-^ mice three days, and in Arf6^-/-^ and VAMP2/3^Δ^ mice seven days, post-wounding (Figure 2J, K). This suggests slower cell migration and proliferation in mice with defects in α-granule cargo packaging and/or platelet endocytosis.

**Figure 2.**
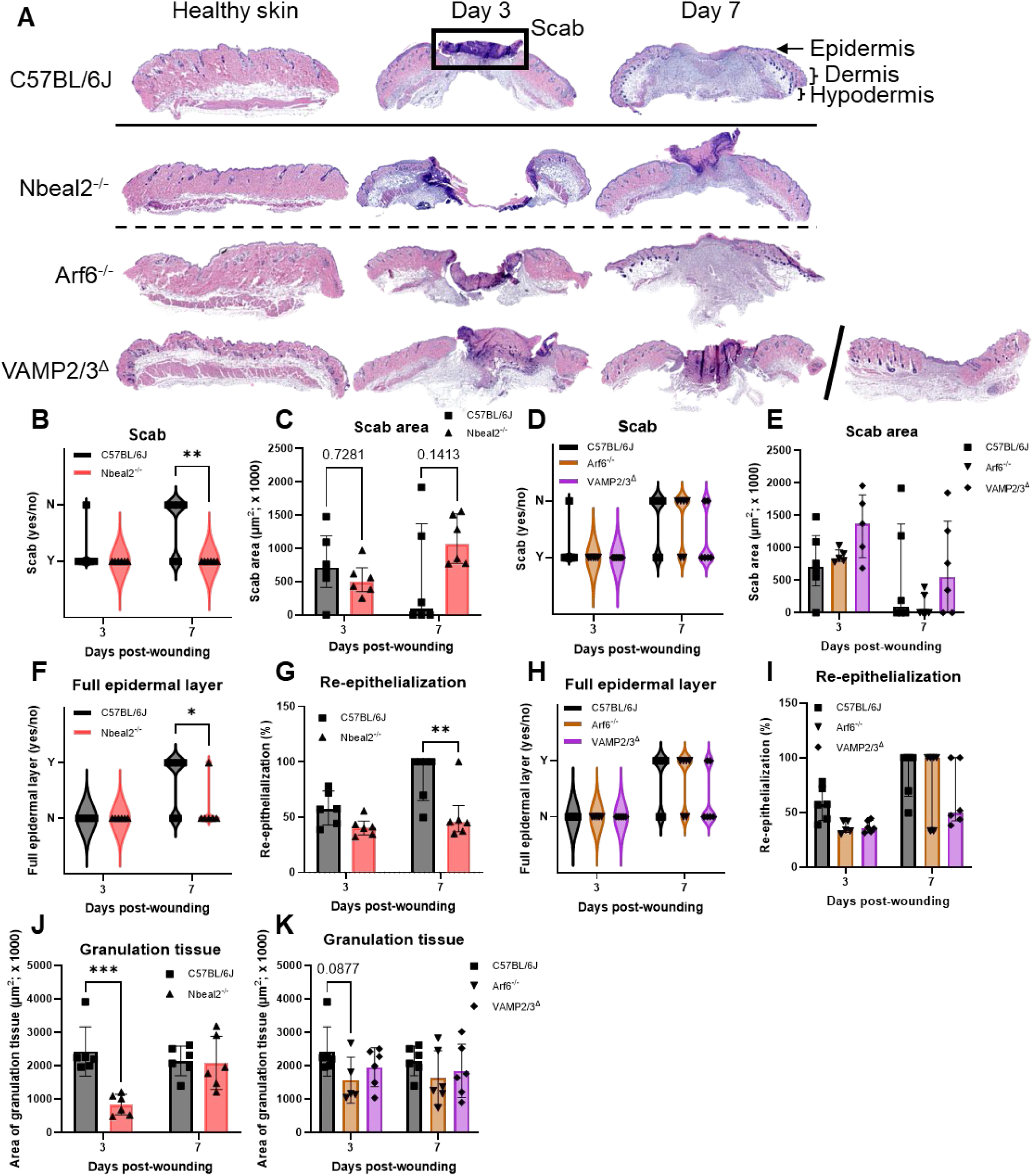
Defective wound healing is reflected by slower scab development and resolution, delayed epidermal regeneration, and slower cell migration and proliferation. **(A)** Representative H&E images of healthy skin and (partially healed) full-thickness wounds on indicated days. Quantification of presence **(B, D)** and area **(C, E)** of a scab, presence of a full epidermal layer **(F, H)**, percentage re-epithelialization **(G, I)**, and granulation tissue area **(J, K)** on day 3 and/or 7 post-wounding in wildtype mice and mice with defects in α-granule cargo packaging **(B, C, F, G, J)** or endocytosis **(D, E, H, I, K)**. Wound scoring: scab absence (N)/presence (Y); full epidermal layer absence (N)/presence (Y). Median ± IQR **(C, E, G, I)**, mean ± SD **(J, K)**, *n* = 5-6, * p < 0.05, ** p < 0.01, *** p < 0.001. Statistics: two-way ANOVA followed by Šidák’s **(B, C, F, G, J)** and Dunnett’s **(D, E, H, I, K)** multiple comparisons tests

Next, structural characteristics of the healthy skin and healing wounds were assessed with Masson’s Trichrome staining, which differentiates between collagen (blue stain) and muscle/keratin (red stain) (Figure 3). In healing wounds, the amount of collagen is indicative of the extent of cell turnover and healing of the dermal skin layer. Muscle and keratin content are signs of inner and outer wound healing, through growth and repair of the *panniculus carnosus* (inside; muscle) and scab formation and resolution (outside; keratin). Collagen content in healthy skin was slightly reduced in Arf6^-/-^ mice (p = 0.07, Figure 3A, D). There was a tendency of reduced collagen and muscle/keratin in the wounds of Nbeal2^-/-^ mice three days post-wounding (p = 0.12 and p = 0.10, respectively; Figure 3A-C), proportionate to a slightly smaller scab size and significant decrease in area of granulation tissue (Figure 2A, C, J). Likewise, wounds of Arf6^-/-^ and VAMP2/3^Δ^ mice had a significant reduction in collagen and muscle/keratin, only seven days post-wounding (Figure 3A, D, E). Thus, wound healing in Nbeal2^-/-^, Arf6^-/-^, and VAMP2/3^Δ^ mice was also structurally disrupted, with potentially specific functions for *de novo* versus endocytosed α-granule cargo release.

**Figure 3.**
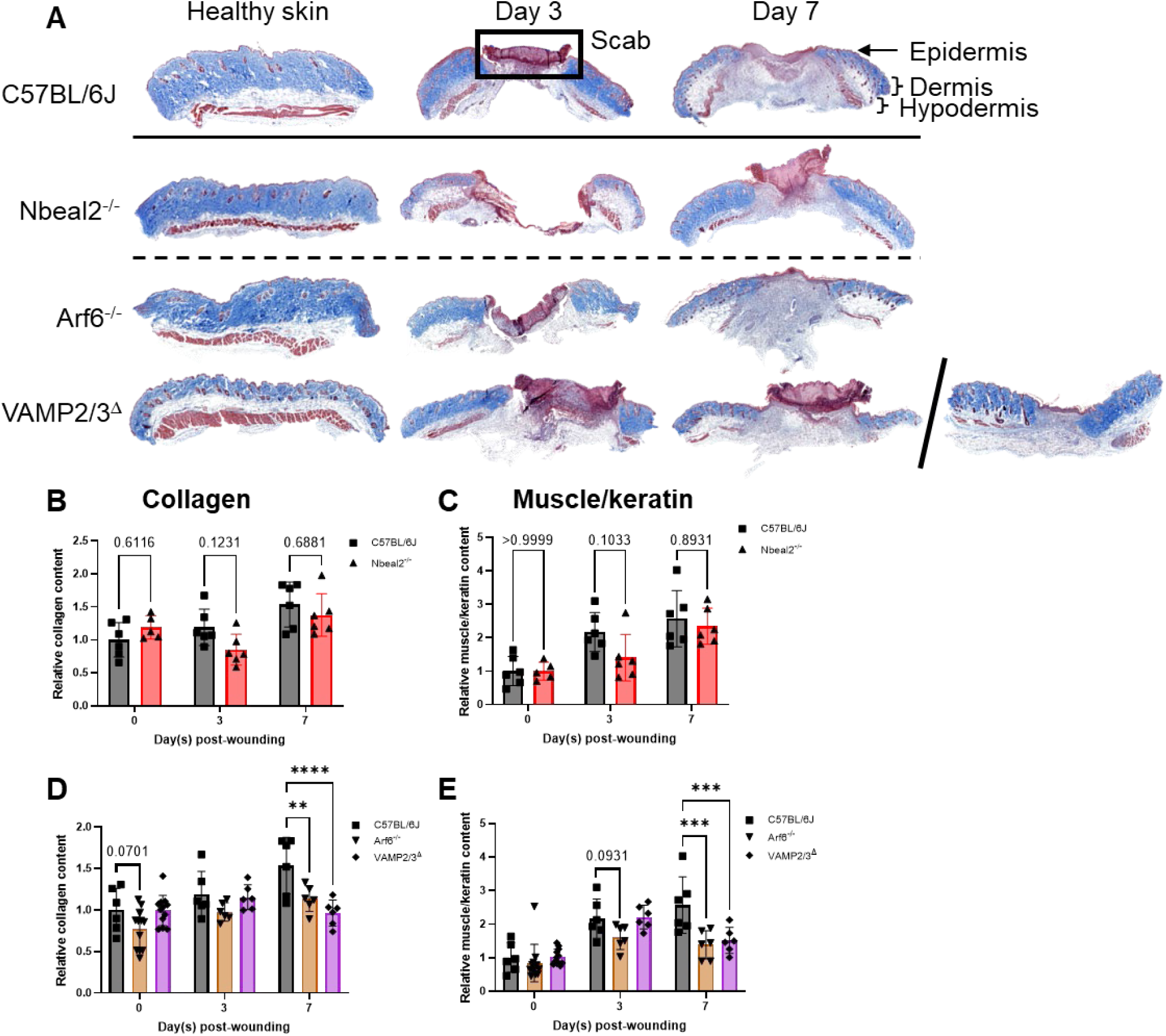
Disrupting α-granule cargo levels and platelet endocytosis changes collagen and muscle/keratin content of healing skin wounds. **(A)** Representative Masson’s Trichrome images of healthy skin and (partially healed) full-thickness wounds on indicated days. Relative collagen **(B, D)** and muscle/keratin **(C, E)** content in wildtype mice and mice with defects in α-granule cargo packaging **(B, C)** or endocytosis **(D, E)**. Mean ± S.D., *n* = 5-12 (day 0), 6 (day 3 and day 7), ** p < 0.01, *** p < 0.001, **** p < 0.0001. Statistics: two-way ANOVA followed by Šidák’s **(B, C)** and Dunnett’s **(D, E)** multiple comparisons tests

### Platelet function in skin wound healing extends to the inflammation, proliferation, and remodeling phases

To gain more mechanistic understanding of platelets’ function in wound healing, we performed an extensive molecular profiling analysis of the skin wound extracts. The panel of analytes was selected based on their presence in platelet releasates and their main function in platelet activation, inflammation, proliferation, and tissue remodeling. A negative correlation between wound resolution and the pro-inflammatory cytokines IL-6 or TNF-α in all mouse strains confirmed that the inflammatory state in the wounds attenuated upon healing (Figure 4A, B, D, E). IL-1β is essential in early wound repair and stimulates inflammatory and proliferative wound healing processes.^64^ However, wounds of Nbeal2^-/-^ and VAMP2/3^Δ^ mice maintained prolonged high levels of IL-1β (Figure 4C, F), perhaps leading to a continuously inflamed environment in the wound environment and contributing to the disrupted healing phenotype in these mice.

**Figure 4.**
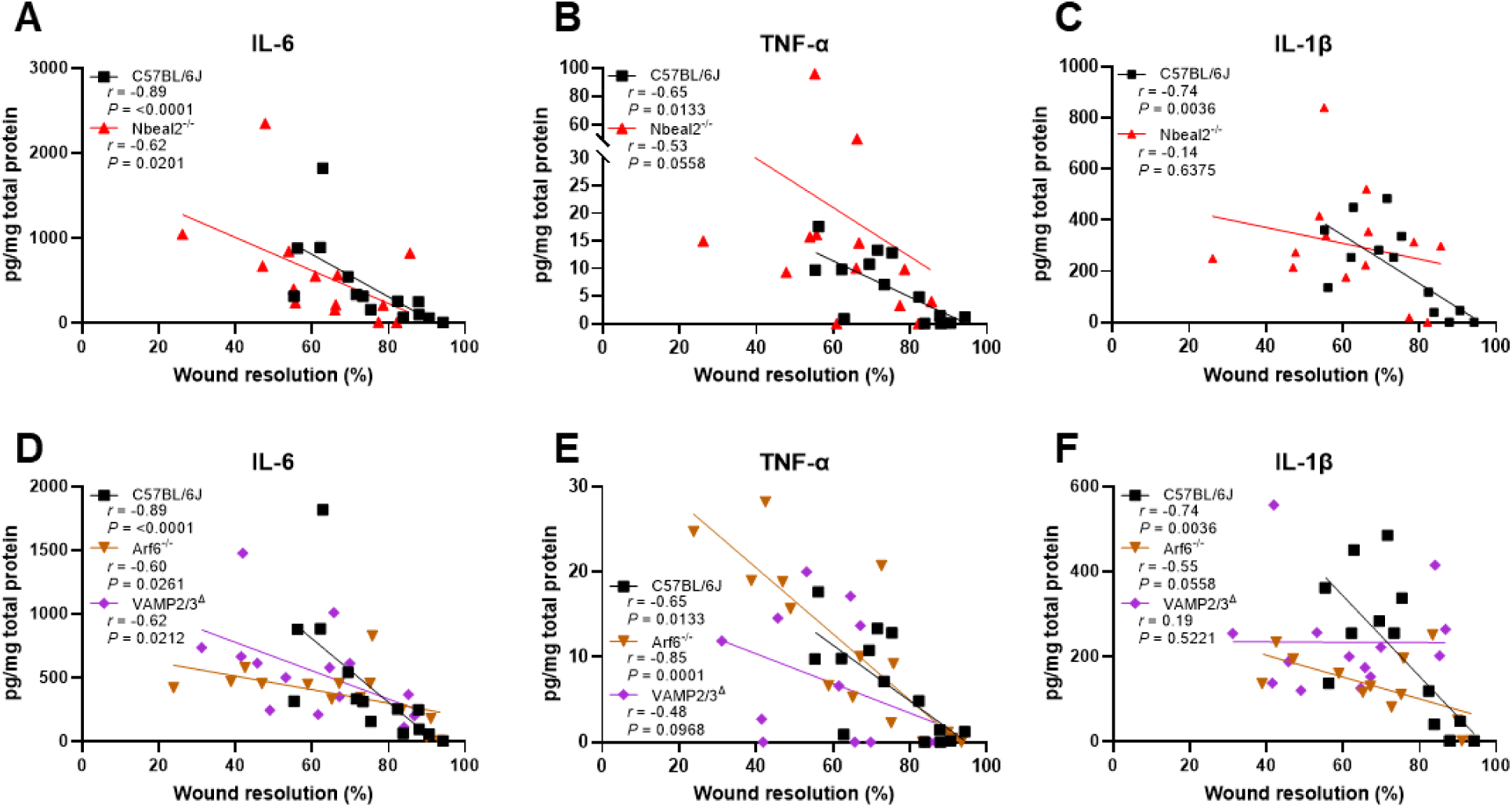
Prolonged high levels of IL-1β suggest a continuously inflamed environment in wounds of Nbeal2^-/-^ and VAMP2/3^Δ^ mice. Spearman correlations between percentage wound resolution and concentrations of IL-6 **(A, D)**, TNF-α **(B, E)**, and IL-1β **(C, F)** in solubilized wound extracts of wildtype mice and mice with defects in α-granule cargo packaging **(A-C)** or endocytosis **(D-F)** after mechanical homogenization. Each datapoint represents one sample. *n* = 13-15

The proliferative phase of wound healing is characterized by the expression of growth factors, including IGF-1, FGF2, and VEGF.^61^ IGF-1 is important for the transition between the inflammatory and proliferative phase through its anti-inflammatory and pro-angiogenic functions and by stimulating keratinocyte migration and proliferation.^65,66^ IGF-1 levels in wounds of wildtype, Nbeal2^-/-^, and VAMP2/3^Δ^ mice reduced during healing (Figure 5A, D). Interestingly, Arf6^-/-^ mice had lower IGF-1 levels in their skin wounds, independent of the percentage wound resolution (Figure 5D). FGF2 regulates wound healing via various cellular processes, including wound re-epithelialization, angiogenesis, and collagen deposition.^65^ The correlations between wound FGF2 levels and wound resolution in Nbeal2^-/-^ mice were in opposite directions compared to wildtype or endocytosis-defective mice (Figure 5B, E). In wildtype mice or mice with platelet endocytosis defects, FGF2 increased as wounds were healing, while in Nbeal2^-/-^ mice, wound FGF2 levels only slightly decreased upon wound resolution. This is in line with the histological data, showing that healing wounds of mice with defective α-granule cargo packaging or platelet endocytosis have reduced re-epithelialization and collagen content (Figures 2G, I, and 3B, D). The effects of FGF2 on angiogenesis might be conveyed in the levels of the pro-angiogenic growth factor VEGF, which is both a *de novo* and endocytosed platelet α-granule cargo protein.^67^ Wounds of wildtype mice showed mildly decreasing VEGF concentrations upon wound resolution, while wound VEGF levels in Nbeal2^-/-^ and Ar6^-/-^ mice were consistently low (Figure 5C, F). Though global VEGF concentrations in VAMP2/3^Δ^ mice were reduced compared to wildtype mice, there was still a negative correlative trend between wound VEGF and wound resolution (Figure 5F).

**Figure 5.**
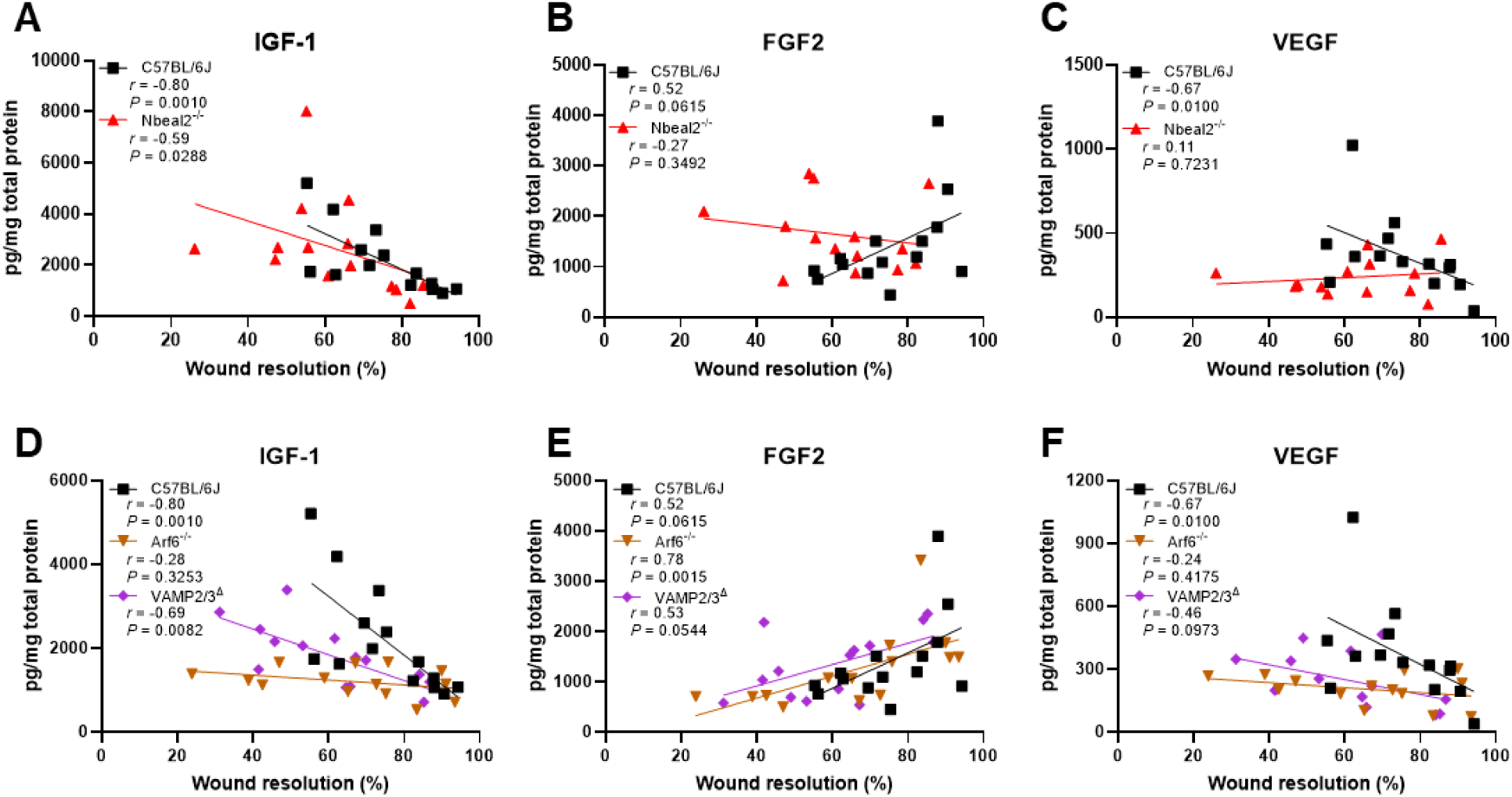
Cell migration, proliferation, and angiogenesis in healing skin wounds requires platelet α-granule cargo release. Spearman correlations between percentage wound resolution and concentrations of IGF-1 **(A, D)**, FGF2 **(B, E)**, and VEGF **(C, F)** in solubilized wound extracts of wildtype mice and mice with defects in α-granule cargo packaging **(A-C)** or endocytosis **(D-F)** after mechanical homogenization. Each datapoint represents one sample. *n* = 13-14

The last phase of skin wound healing is maturation, or the tissue remodeling phase. Wildtype mice showed clear correlations between wound resolution and the expression of tissue remodeling factors, *i.e.,* MMP-3, MMP-9, and TIMP-1, in the wound extracts (Figure 6A-F). In wildtype mice, MMP-3 concentrations were high when wounds were healed to about 50-75%, after which they quickly dropped (Figure 6A, D). MMP-3 stimulates activation of MMP-9,^68^ and consistently, the levels of MMP-9 in the wound extracts followed an inverse pattern compared to MMP-3 (Figure 6B, E). Tissue remodeling in wildtype mice further correlated with a concurrent decrease of TIMP-1 in extracts of healing wounds (Figure 6C, F). The direction and slope of these factors did not correlate with healing in Nbeal2^-/-^ mice, indicating disrupted tissue remodeling (Figure 6A-C). Arf6^-/-^ and VAMP2/3^Δ^ mice exhibited more complex patterns (Figure 6D-F). Here, no clear correlation existed between MMP-3 concentrations and wound resolution, as also seen in Nbeal2^-/-^ mice (Figure 6D). Overall MMP-9 levels were extremely low, almost absent, until the last phases of wound repair, when they acutely increased (Figure 5E). Like wound VEGF concentrations (Figure 5F), TIMP-1 expression was lower in skin wound extracts of Arf6^-/-^ and VAMP2/3^Δ^ mice than in wildtype mice. However, there was still a clear decrease in wound TIMP-1 levels that significantly inversely correlated with (an increase in) wound resolution (Figure 6F).

**Figure 6.**
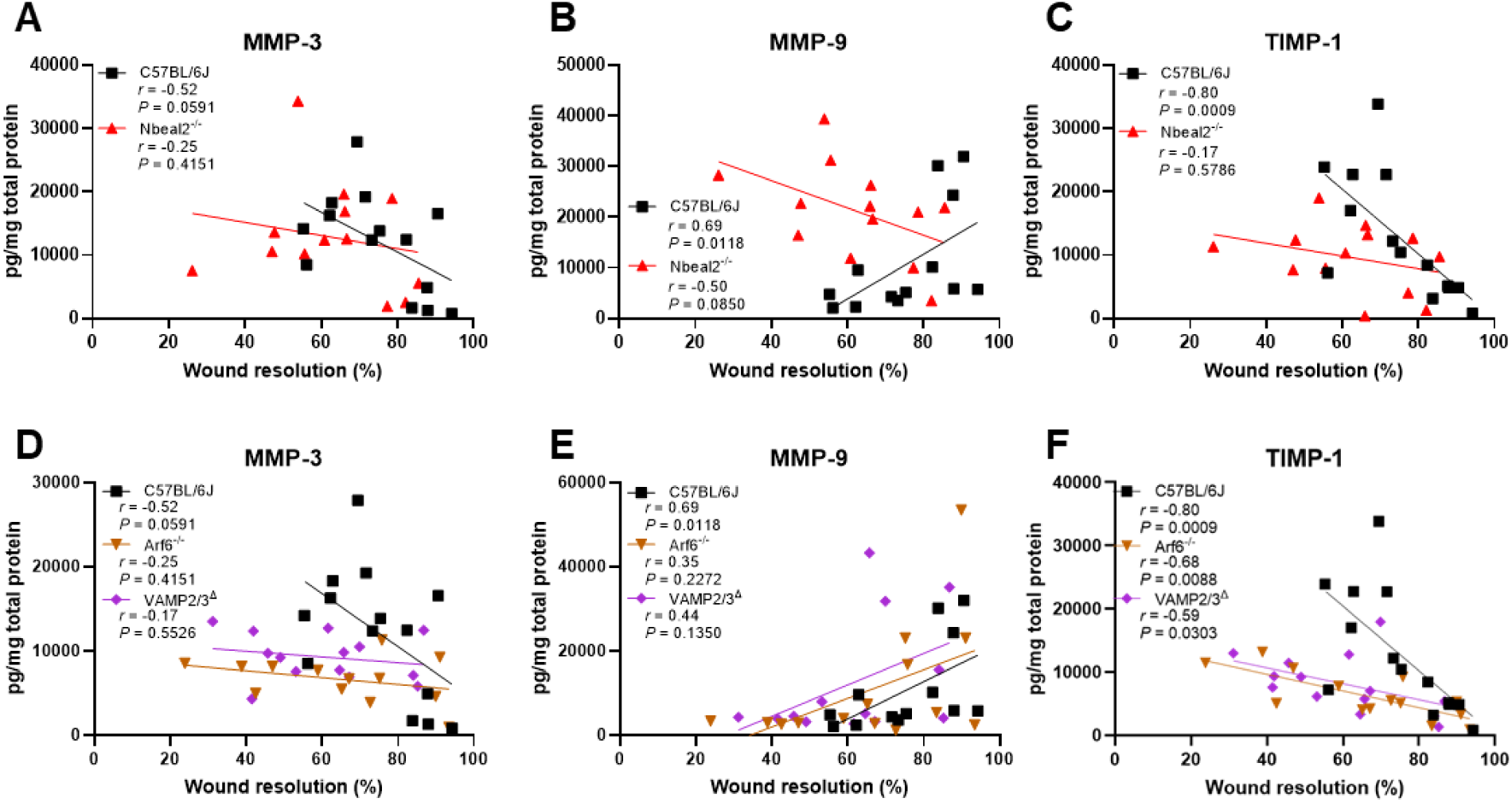
Disproportional correlations of MMP-3, MMP-9, and TIMP-1 levels in healing wounds of Nbeal2^-/-^, Arf6^-/-^, and VAMP2/3^Δ^ mice indicate disturbed tissue remodeling. Spearman correlations between percentage wound resolution and concentrations of MMP-3 (A, D), MMP-9 (B, E), and TIMP-1 (C, F) in solubilized wound extracts of wildtype mice and mice with defects in α-granule cargo packaging (A-C) or endocytosis (D-F) after mechanical homogenization. Each datapoint represents one sample. *n* = 13-14

## Discussion

This study is the first to use specific transgenic mouse models with defined platelet defects to address how platelet α-granule cargo affects physiological (skin) wound healing. We demonstrated that proper healing of dorsal skin wounds, in mice, requires release of both *de novo* and endocytosed platelet α-granule cargo. Notably, platelet function does not appear to be limited to the hemostasis phase of wound healing; instead, it expands to all phases, affecting the presence and levels of various bioactive molecules in the skin microenvironment via their endo- and exocytic pathways. While these alterations in the expression of bioactive molecules in skin wounds could be directly due to platelet releasate, it is also possible that platelets stimulate the production of such molecules by other cell types. However, it should be noted that our data confirms a role for platelets in these processes since the Arf6^-/-^ and VAMP2/3^Δ^ are platelet-specific deletions.

Interest in and research on the role of platelets and coagulation in wound healing has increased over the past decades. However, it has been mainly focused on the effects of exogenous application of platelet-rich plasma or platelet-derived growth factors to treat chronic, non-healing wounds in pathological conditions, *e.g.,* diabetes.^69–75^ Studies on fundamental endogenous mechanisms of platelet function or coagulation in physiological, or acute, wound repair are still limited and with somewhat contrasting results.^45–47,76–82^ These studies point to a central role for platelets and coagulation in wound healing by using mouse models and inhibitors that target specific platelet receptors or coagulation factors. Building on this and on the clinical use of platelet-rich plasma and platelet-derived growth factors in facilitating wound repair, we more specifically investigated different mechanistic and molecular facets of platelet α-granule cargo in wound healing. Our findings show that inaccurate α-granule cargo packaging or platelet endocytosis led to severely impaired healing kinetics and distinctive wound morphology. A potential key player herein might be fibrinogen. Fibrinogen-deficient mice have severe healing defects of both incisional and excisional wounds.^77^ Moreover, Wichaiyo and colleagues found that concurrent deletion of platelet CLEC-2 and GPVI or dasatinib treatment accelerated wound healing by disrupted vascular integrity, and subsequently increased minor bleeding and fibrin(ogen) deposition into the wound surroundings.^45–47^ In contrast, wound healing was delayed in mice lacking FIX or with low tissue factor or low FVII, along with increased presence of wound hematomas, but reduced fibrin in the wound.^78,79,82^ This suggests a tight balance between vascular integrity and microbleeds in wound healing, with a potential bilateral role for fibrin(ogen).

We have previously shown that Arf6^-/-^ and VAMP2/3^Δ^ mice have impaired fibrinogen uptake and storage without affecting hemostasis or thrombosis.^51,52^ Platelets of Nbeal2^-/-^ mice also have extremely reduced fibrinogen content.^44^ Nbeal2^-/-^ mice have severely defective hemostasis and arterial thrombosis; however, gray platelet syndrome (GPS) patients with confirmed mutations in *NBEAL2* mostly presented with a mild to moderate bleeding phenotype, though severe bleeding has been reported.^44,54,83^ It is not known whether GPS patients have wound healing complications. Wounds from Arf6^-/-^ and VAMP2/3^Δ^ mice had a similar overall histological appearance as wildtype mice three days after wounding, but (epi)dermal healing lagged in more advanced stages. This was verified quantitatively with a reduction in wound collagen and muscle/keratin content, and slower scab resolution, re-epithelialization, and cell migration and proliferation. Likewise, (epi)dermal regeneration was severely impaired in Nbeal2^-/-^ mice, which was particularly evident in earlier healing stages, with delayed scab formation, re-epithelialization, and cell migration. Consistently, other studies have also shown that acceleration or delay of wound healing after manipulating platelet function can be partly attributed to the amount of re-epithelialization and granulation tissue formation.^44,45,47^ Interestingly, Deppermann *et al.* showed that epithelial wound closure in Nbeal2^-/-^ mice was similar as in wildtype mice seven days post-wounding, though (collagenous) granulation tissue and myofibroblasts were reduced.^44^

Aside from fibrin(ogen), disrupted vascular integrity potentially allows for distribution of other bioactive molecules into the injured area, such as cytokines and growth factors. Using an extensive molecular profiling analysis of solubilized skin wound extracts, we found that expression of various proteins was altered in healing skin wounds of Nbeal2^-/-^, Arf6^-/-^, and VAMP2/3^Δ^ mice. These expression patterns affected the complete wound healing process, lasting to at least seven days after wounding. We categorized the measured analytes based on their main function to more specifically study the inflammation, proliferation, and remodeling phases of wound healing. The pro-inflammatory cytokine IL-1β plays a crucial role in early stages of wound repair by stimulating the immune response, fibroblast proliferation, and angiogenesis.^64^ However, prolonged IL-1β activity is detrimental to the wound healing process and its expression has been shown to be a key regulator in (dysfunctional) healing of diabetic wounds.^64,84–87^ Chronic wounds, such as observed in diabetic patients, are associated with infection and an excessive and prolonged inflammatory state with increased levels of dysfunctional neutrophils, macrophages, and other immune cells.^27^ We found constant high levels of IL-1β in skin wound extracts of Nbeal2^-/-^ and VAMP2/3^Δ^ mice, independent of the extent of wound resolution. It should be noted that the various wound healing phases overlap; the presence and activity of certain proteins often regulates the expression of others. As such, IL-1β coordinates tissue remodeling via the expression of matrix metalloproteinases (MMPs), including MMP-9.^84^ Re-epithelialization and matrix remodeling processes are also disturbed in chronic wounds, partly due to increased protease levels and associated degradation of growth factors and cytokines.^27^ In this study, Nbeal2^-/-^ mice, and to a slightly different extent Arf6^-/-^ and VAMP2/3^Δ^ mice, had extremely dysregulated tissue remodeling, which was apparent in the expression of MMP-3, MMP-9, and TIMP-1 in wound extracts and their correlation with wound resolution. The inflammation and tissue remodeling wound healing phases are connected through the proliferation phase. TGF-β is a multifunctional cytokine involved in both proliferation and tissue remodeling and recent studies have found a platelet-mediated role for TGF-β in skin wound healing.^44,64,70^ Angiogenesis is a central process during proliferative wound repair, with an important function for VEGF. VEGF is expressed in a broad range of cells and, in platelets, released from α-granules, where it is packaged both via the *de novo* as well as the endocytosis pathway.^67,88–90^ VEGF levels were consistently low in skin wound extracts of Nbeal2^-/-^ and Arf6^-/-^ mice, indicating insufficient angiogenesis.

It has been shown that, in vascular inflammation, platelets can migrate and act as mechano-scavengers to clear bacteria from the environment.^91–94^ By doing so, platelets could potentially use their migratory capabilities and endocytic trafficking machinery to clear the injury site from pathogens or inflammatory stimuli, such as Pathogen- or Damage-Associated Molecular Patterns (PAMPs or DAMPs). We have previously demonstrated that the uptake of HIV-1 pseudovirions was decreased in platelets from VAMP3^-/-^ and Arf6^-/-^ mice.^95^ In this study, comparisons between Arf6^-/-^ and VAMP2/3^Δ^ mice demonstrated differences in skin wound healing responses and thus highlight important aspects of endocytosis signaling in wound repair.

Altogether, this study shows that both *de novo* and endocytosed cargo proteins from platelet α-granules are essential for skin wound healing, even beyond the initial hemostasis phase. This research will improve clinical applications directed towards advanced treatment of acute and chronic wounds. Recent fundamental and clinical studies investigated topical or intravenous application of platelet-derived factors in diseased conditions.^70,73,96,97^ However, more extensive research, such as transcriptomics and proteomics, should be performed to gain insight into which cargo proteins are most essential for skin wound healing, particularly addressing the differences between *de novo* and endocytosed platelet α-granule cargo. Furthermore, additional studies should focus on the temporal roles of platelets and their interaction with the skin microenvironment.

## Acknowledgements

The authors are thankful for the efforts of Ming Zhang in managing the mouse colony. The authors thank the University of Kentucky Pathology Research Core and Light Microscopy Core for their technical and experimental assistance. The work was supported by grants from the American Heart Association (23POST1020159) to D.M.C., from the NIH/NHLBI (R35HL150818) to S.W.W., a Department of Veterans Affairs Merit Award to S.W.W, and an Institutional Development Award from the NIH/NIGMS (P30GM127211).

## Author contributions

D.M.C. conceived the project, performed and analyzed the experiments, and wrote the manuscript. H.R.A. assisted with some of the experiments. S.W.W. directed the research and edited the manuscript.

