## Supplemental material for "Platelet endocytosis and α-granule cargo packaging are essential for normal skin wound healing"

### Supplemental Methods

#### Genotyping

The genotyping primers and conditions were as follows for the Nbeal2$^{-/-}$ mice (RRID:MMRRC_031611-UCD):

- Common primer 5'-GTCCTGCTTGACCTACCGTC;
- Wildtype primer 5'-CAGGGAGGATAACGAGATAGTCTT;
- Knockout primer 5'-CCTAGGAATGCTCGTCAAGA.

The PCR conditions were 95 °C, 30 sec; 64 °C, 30 sec; 70 °C, 1 min; repeated for 40 cycles.

The genotyping primers and conditions were as follows for the Arf6$^{-/-}$ mice:

- Floxed Arf6 gene, forward primer 5′-GACCCCATGAGTGTTGTCAC;
- Floxed Arf6 gene, reverse primer 5′-GGGATACATAGAGAAACCTTGTCTCAGG.

The PCR conditions for the floxed Arf6 gene were 94 °C, 10 minutes for 1 cycle; 94 °C, 1 min; 54 °C, 45 sec; 72 °C, 30 sec; repeated for 30 cycles; 72 °C, 7 min.

- PF4-Cre transgene, forward primer 5'-CCCATACAGCACACCTTTTG;
- PF4-Cre transgene, reverse primer 5'-TGCACAGTCAGCAGGTT.

The PCR conditions for the PF4-Cre transgene were 94 °C, 7 minutes for 1 cycle; 94 °C, 1 min; 50 °C, 1 min; 72 °C, 1.5 min; repeated for 30 cycles; 72 °C, 10 min.

The genotyping primers and conditions were as follows for the VAMP2/3$^{\Delta}$ mice:

- RC::PFtox, forward primer 5'-GCCGATCACCATCAACAACTTC;
- RC::PFtox, reverse primer 5'-GCAGAGCTTCACCAGCAACG.

The PCR conditions for the RC::PFtox were 94 °C, 4 minutes for 1 cycle; 94 °C, 30 sec; 58 °C, 30 sec; 72 °C, 1 min; repeated for 35 cycles; 72 °C, 5 min.

- PF4-Cre transgene, forward primer 5'-CCCATACAGCACACCTTTTG;
- PF4-Cre transgene, reverse primer 5'-TGCACAGTCAGCAGGTT.

The PCR conditions for the PF4-Cre transgene were 94 °C, 7 minutes for 1 cycle; 94 °C, 1 min; 50 °C, 1 min; 72 °C, 1.5 min; repeated for 30 cycles; 72 °C, 10 min.

#### Supplemental Table

**Supplemental Table 1. Analytes measured in the mouse premixed multi-analyte Luminex Discovery Assays and their transcriptomic and proteomic expression in human megakaryocytes (MKs) and platelets**

| Abbreviation | Full name | MK RNA | Platelet RNA | Platelet protein |
|---|---|---|---|---|
| **Platelet activation markers** | | | | |
| CCL5/RANTES | CC chemokine ligand 5/regulated upon activation, normal T cell expressed and secreted | Yes | Yes | Yes |
| CD62P | P-selectin | Yes | Yes | Yes |
| **Inflammatory markers** | | | | |
| IL-1β | Interleukin-1β | Yes | Yes | No |
| IL-6 | Interleukin-6 | No | Yes | No |
| IL-10 | Interleukin-10 | N.I. | N.I. | N.I. |
| IL-12 | Interleukin-12 | No | Yes | No |
| TNF-α | Tumor necrosis factor-α | Yes | Yes | No |
| **Growth factors** | | | | |
| EGF | Epidermal growth factor | Yes | Yes | Yes |
| FGF2 | Fibroblast growth factor 2 | Yes | Yes | No |
| IGF-1 | Insulin-like growth factor-1 | No | Yes | No |
| PDGF-AA | Platelet-derived growth factor AA | Yes | Yes | Yes |
| PDGF-BB | Platelet-derived growth factor BB | Yes | Yes | Yes |
| VEGF | Vascular endothelial growth factor A | Yes | Yes | No |
| | Vascular endothelial growth factor B | Yes | Yes | No |
| | Vascular endothelial growth factor C | Yes | Yes | Yes |
| **Tissue remodeling factors** | | | | |
| MMP-2 | Matrix metalloproteinase-2 | N.I. | N.I. | N.I. |
| MMP-3 | Matrix metalloproteinase-3 | N.I. | N.I. | N.I. |
| MMP-9 | Matrix metalloproteinase-9 | Yes | Yes | Yes |
| TIMP-1 | Tissue inhibitor of metalloproteinases-1 | Yes | Yes | Yes |
| TIMP-4 | Tissue inhibitor of metalloproteinases-4 | No | Yes | No |

Analytes are subdivided into platelet activation markers, inflammatory markers, growth factors, and tissue remodeling factors. MK, megakaryocyte; N.I, not identified. Transcriptomic and proteomic data derived from: Huang J *et al.*, Sci Rep, 2021.[1]

### Supplemental Figure

#### Supplemental Figure 1

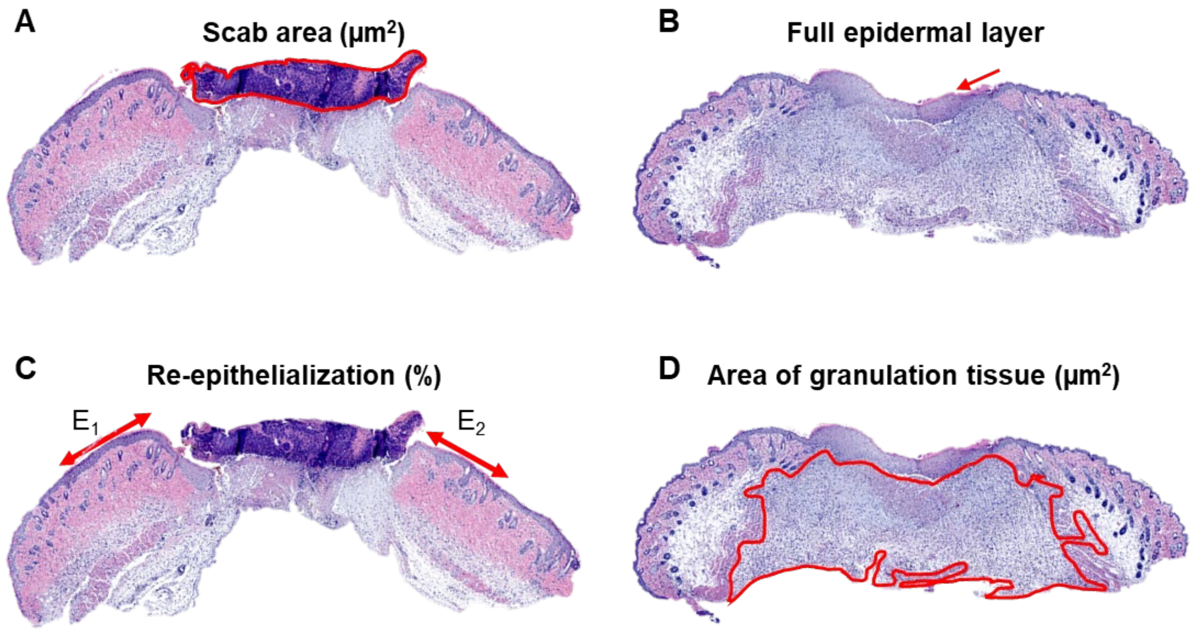


**Supplemental Figure 1. Representative H&E images and analysis examples of several wound morphometrics**, *i.e.,* scab presence and area **(A)**, presence of a full **(B)** or partial **(C)** epidermal layer, percentage re-epithelialization **(C)**, and area of granulation tissue **(D)**.
